## Supplementary figures and images for "Noise leads to the perceived increase in evolutionary rates over short time scales"

### Movie S1

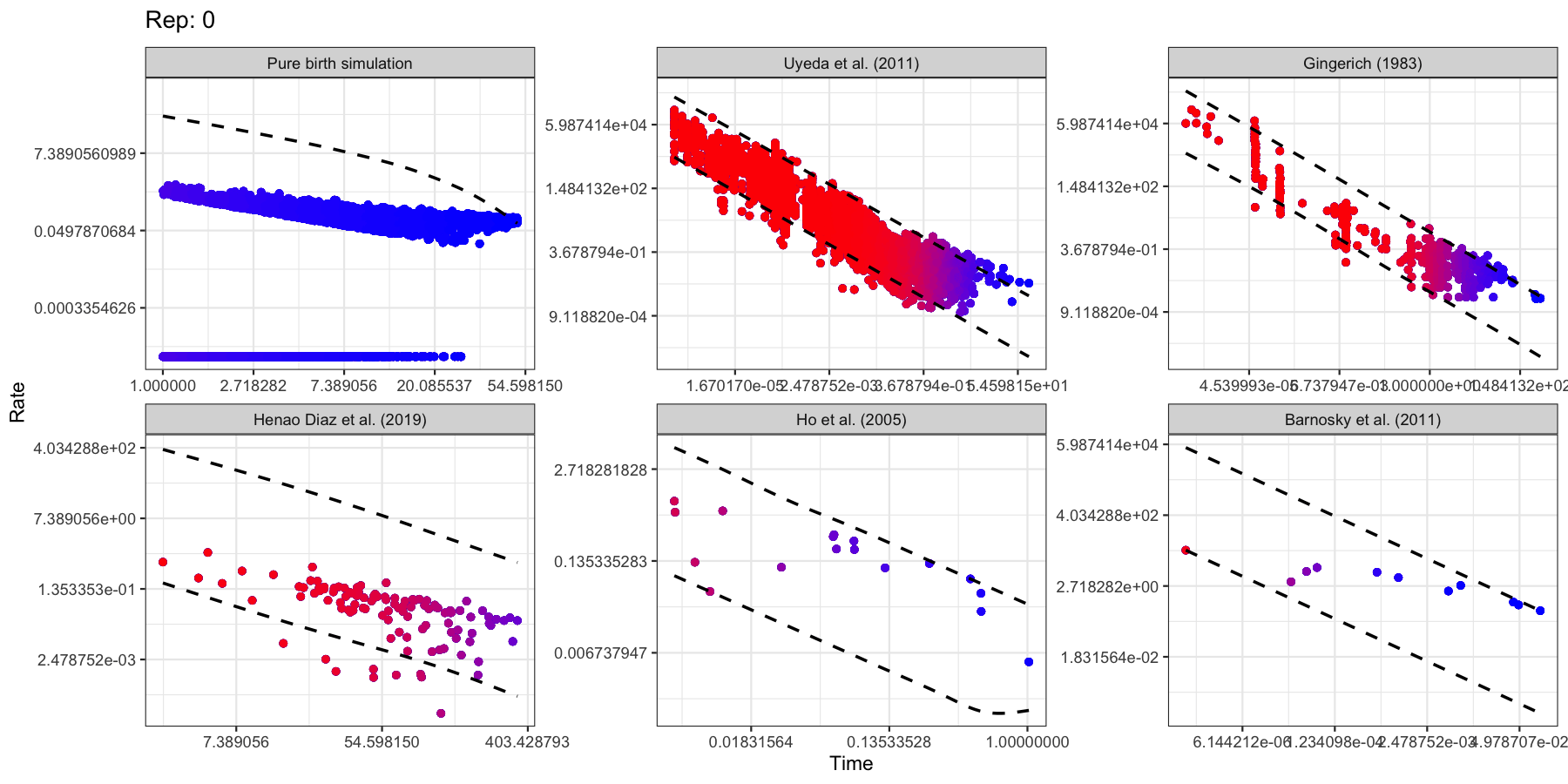
